# Identification and Classification of Fungal GPCR Gene Families

**DOI:** 10.1101/2024.11.25.625238

**Authors:** Zhiyin Liu, Asaf Salamov, Igor V. Grigoriev

**Affiliations:** U.S. Department of Energy Joint Genome Institute, Lawrence Berkeley National Laboratory, Berkeley, CA 94720, USA; Department of Plant and Microbial Biology, University of California Berkeley, Berkeley, CA 94720, USA; Division of Life Science, The Hong Kong University of Science and Technology, Hong Kong SAR, China

## Abstract

G protein-coupled receptors (GPCRs) are transmembrane proteins crucial for signal transduction in eukaryotes, responding to diverse extracellular signals. Researchers have found 14 distinct types of GPCRs in fungi but their distribution among numerous fungal species remained largely unexamined. Our study identified and classified GPCRs in 1,357 fungal species, and shed light on GPCR distribution in fungi. The predominant class detected in fungi was Pth11-like GPCRs, exclusively found in Pezizomycotina and are notably acknowledged for their involvement in fungal pathogenicity. Our analysis suggested that Pezizomycotina ancestor possessed a more extensive array of Pth11-like GPCRs, but over time, some species underwent considerable reductions in these GPCRs in conjunction with genome contractions. Additionally, we identified 2,089 mammalian homologs in Rhodopsin, Glutamate, and Frizzled classes across 594 fungal species, thereby augmenting the recognized fungal GPCR classes by three classes thought to be mammalian-specific. Utilizing a custom-built convolutional neural network (CNN) for the identification of fungal GPCRs, we discovered several potential novel fungal GPCRs. Moreover, anticipated interactions between these prospective new GPCRs and G-alpha proteins, as simulated by AlphaFold Multimer, offered further confirmation for these findings.

## Introduction

G protein-coupled receptors (GPCRs) is a large and multifaceted family of membrane-bound signaling proteins with seven TMHs. These receptors in eukaryotes mediate a plethora of extracellular inputs into the cell, which in turn regulate several biological processes including but not limited to vision, taste, neuroendocrine functions and immunity [1, 2]. GPCRs are essential to the survival and adaptation of fungi, which are largely utilized in sensing the environment, sexual development, pathogenesis and many other functions [3–6]. These receptors enable the perception of external cues, which is essential for fungi as they navigate their varied habitats, often characterized by nutrient scarcity and environmental adversities [7, 8]. Despite the extensive characterization of GPCRs in mammals, their counterparts in fungi remain comparatively underexplored and only a fraction of mammal diversity has been mirrored in fungal species [3].

To date, fungal GPCRs have been classified into 14 distinct classes [9]. Among these, six are well-established classical GPCRs. The first five classes are Ste2 pheromone receptors [10], Ste3 pheromone receptors [10], glucose-sensing Gpr1 homologs [11, 12], nutrient sensors analogous to Stm1 in *Schizosaccharomyces pombe* [13], and cAMP receptors homologous to those in *Dic-tyostelium discoideum* [14]. These receptors activate two key signaling pathways, the mitogen-activated protein kinase (MAPK) cascades and the cAMP-dependent protein kinase A (PKA) pathway, which are essential for fungal physiology [15]. The sixth classical class (GPCR Class 9) is microbial opsins [16]. In addition, eight novel fungal GPCR classes 6–8 [17–19] and classes 10–14 [18, 20–23] have also been identified. These include GprK-like receptors with RGS domains [17], receptors similar to the rat growth hormone-releasing factor receptor [18], mPR-like/PAQR receptors [19], the Lung 7TM superfamily [20], GPCR89/ABA receptors [21], Family C-like receptors [22], the DUF300 superfamily/PsGPR11 [23], and Pth11-like receptors [18]. However, the current understanding of fungal GPCRs is restricted to a small number of species. This study investigates the distribution of all 14 fungal GPCR classes that have been previously described and three additional classes that have been previously described exclusively in mammals across a diverse array of fungal species, addressing the stark disparity between the extensive research on mammalian GPCRs and the nascent understanding of their fungal counterparts.

According to GRAFS GPCR classification scheme, there are 6 families of mammalian GPCRs: Glutamate, Rhodopsin, Adhesion, Frizzled, Taste2, and Secretin [24]. However, the existing categorization of fungal GPCRs does not yet incorporate these families, and only Class 7 includes the Pfam domain 7tm 2 homologous to the mammalian Secretin GPCR family. This presents us an opportunity to investigate potential similarities between fungal and mammalian GPCRs and to expand the number of potential mammalian homologous classes in fungi.

Among the several types of GPCRs, Pth11-like GPCR has been found to be a major determinant in late endosomal homotypic fusion and protein sorting in *Magnaporthe oryzae*, thereby influencing appressorium growth, cAMP generation, and finally contributing to the pathogen’s virulence [25]. Notably, previous studies show that Pth11-related GPCRs were only found in fungi belonging to Pezizomycotina, while none were found in other subphyla of Ascomycota or Basidiomycota [18, 25, 26]. Pth11 contains an amino-terminal extracellular cysteine-rich CFEM domain (pfam05730), which is involved in diverse fungal processes, including pathogenesis in *Magnaporthe grisea* and *Candida albicans*, as well as cell wall biogenesis and stability in non-pathogenic fungi such as *Saccharomyces cerevisiae*. However, the CFEM domain is present in only a small number of Pth11-like proteins from *M. grisea* and *Neurospora crassa* [27]. The abundance of Pth11-like GPCRs across species remains unknown, despite their extensive distribution in Pezizomycotina.

Due to the varied functions and evolutionary distinctions of GPCRs, forecasting their presence and roles in different fungal species is exceedingly difficult. Recent improvements in deep learning, particularly in protein function prediction, present new opportunities. Recent advancements in natural language processing and sequential data preprocessing have highlighted the utilization of deep neural networks (DNN), convolutional neural networks (CNN), recurrent neural networks, long short-term memory networks, and attention-based transformer models [28]. Furthermore, features that are derived from amino acid sequences and three-dimensional (3D) protein structures can be incorporated into multi-modal deep learning techniques. Approximately half of protein function prediction tools employ 1D CNNs combined with DNNs, demonstrating efficacy in sequence classification tasks [29], including ProteInfer [30] and DeepFRI [31]. ProteInfer utilizes deep CNN to directly forecast Enzyme Commission numbers and Gene Ontology terms from unaligned amino acid sequences. Conversely, DeepFRI employs a graph convolutional network that integrates characteristics from a protein language model and structural data.

A key criterion for identifying GPCRs is their ability to activate G-proteins. Machine learning-based protein-protein interaction (PPI) models, such as those using natural language processing techniques (e.g., D-script [32]), are increasingly used to predict these interactions alongside traditional wet lab methods. Advances in 3D structure prediction, especially via AlphaFold Multimer [33], has improved prediction of interactions within protein complexes.

Here, we performed an extensive examination of 1,357 fungal species, finding GPCRs to illustrate a varied range of GPCRs and highlight the variations in their distributions among species. Additionally, we examined the distribution of Pth11-like GPCRs in Pezizomycotina and the evolutionary dynamics of these receptors among closely related species within the class Sordariomycetes. We also investigated possible relationships between Pth11-like GPCRs and host-pathogen interactions as well as secondary metabolism. Following the identification of known GPCRs among different fungal species, we predicted novel GPCRs featuring seven transmembrane helices (TMHs) using a one-dimensional CNN binary classifier specifically designed for the identification of fungal GPCRs. We further integrated predictions from DeepFRI and ProteInfer, fol-lowed by modeling interactions between candidate GPCRs and G-alpha proteins using AlphaFold Multimer to identify high-confidence GPCR-G-alpha combinations for further investigation.

## Results

### Distribution of known fungal GPCR classes in 1,357 fungal species

Using 67 literature-derived GPCRs from 14 known fungal GPCR classes (Supplementary file 1 Table S1) queried against 1,357 fungal genomes from the MycoCosm database, we found 31,493 potential GPCR sequences across three fungal groups: Ascomycota, Basidiomycota, and the Early Diverging Fungi (EDF). Significantly, no GPCRs were detected in 25 species (Supplementary file 1 Table S2). Ascomycota demonstrated the greatest average abundance and diversity among species and GPCR classes, accounting for 75.9% of predicted fungal GPCRs. Conversely, species within Basidiomycota and EDF exhibited many absent or reduced GPCRs at both the species and GPCR class levels in comparison to Ascomycota (Fig. 1a).

**Figure 1:**
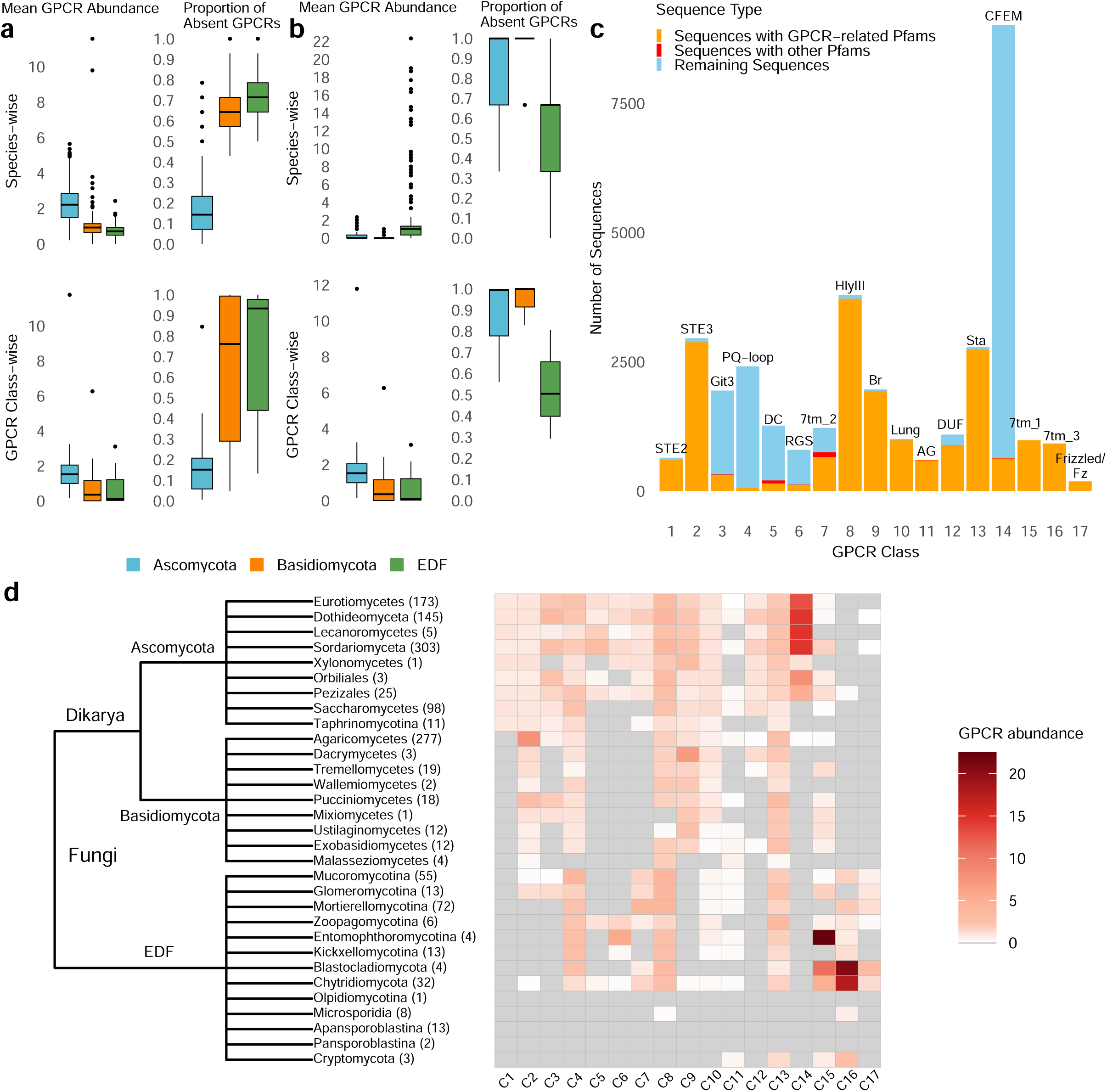
GPCR distribution among 1,357 fungal species. **a** Box plots showing the mean GPCR abundance and proportion of absent GPCRs across fungi and 14 known fungal GPCR classes. **b** Box plots demonstrating the mean GPCR abundance and proportion of absent GPCRs across fungi and 3 mammalian GPCR classes. The settings of the figure are the same as in **a**. **c** Pfam domain distribution across all the 17 GPCR classes. The x-axis represents each GPCR class, while the y-axis shows the number of sequences in each class. Bars are color-coded: the orange section indicates sequences containing the specified GPCR-related Pfam domain (displayed above each bar), the red section represents sequences with other non-GPCR Pfam domains, and the blue section represents sequences without Pfam domains. DC in Class 5 refers to Dicty_CAR, Br in Class 9 refers to Bac_rhodopsin, Lung in Class 10 refers to Lung_7-TM_R, AG in Class 11 refers to ABA_GPCR, DUF in Class 12 refers to DUF3112, and Sta in Class 13 refers to Solute_trans_a. **d** Distribution of GPCRs across 31 fungal clades within three groups. The tree illustrates the taxonomic cladogram of fungal classes. The adjacent heatmap displays the average GPCR abundance for species in each fungal class. Numbers following the fungal class names at the tree nodes indicate the number of species within each taxonomic group.

Further classification of the fungal species into 31 clades showed that species from three groups, Olpidiomycotina, Apansporoblastina, and Pansporoblastina, completely lacked GPCRs across all 14 recognized categories (Fig. 1d). Olpidiomycotina is a subphylum comprising a category of obligate parasitic fungi. Apansporoblastina and Pansporoblastina are classified within the phylum Microsporidia, comprising spore-producing unicellular parasites. Furthermore, Class 1 GPCRs, represented by STE2-like fungal pheromone receptors, are exclusively found in Ascomycota, whereas Class 2 GPCRs, which include STE3-like fungal pheromone receptors, are predominantly present in both Ascomycota and Basidiomycota. To be specific, Agaricomycetes in Basidiomycota exhibited the highest abundance of Class 2 GPCRs compared to other GPCR classes and fungal groups. Class 5 (cAMP receptor-like), Class 6 (GprK-like receptors with RGS domains), and Class 7 (rat growth hormone-releasing factor receptor homologs) are exclusively found in Ascomycota and EDF, but are absent in Basidiomycota. Furthermore, Dacrymycetes were characterized by an expansion of Class 9 GPCRs, rhodopsin-like receptors predominant in Ascomycota and Basidiomycota. Pth11-related GPCRs (Class 14), exclusively present in the Pezizomycotina subphylum of Ascomycota, are the most abundant class (28.6%) across all fungi (Fig. 1d). This finding expands on previous studies that identified Pth11-related GPCRs as unique to 20 species of Eurotiomycetes and Sordariomycetes in Pezizomycotina [27], to 659 species, now including species from Dothideomycetes, Lecanoromycetes, Xylonomycetes, Orbiliales, and Pezizales.

Pfam domain characterization was used for additional validation of these sequences, where 16,240 out of 31,493 sequences contained specific domains, with 16,055 (98.9%) of them possessing functional GPCR domains that were consistent with their associated GPCR class (Table 1). While classes 4 and 6 showed the smallest number of sequences with their respective domains, PQ-loop and RGS, Class 8 GPCRs displayed the highest number of sequences including GPCR-related Pfam domain (HlyIII). Furthermore, although Class 14 GPCRs were the most plentiful among fungal speceis, only 6.9% of these sequences included the CFEM domain, which is linked to fungal pathogenicity in Pth11 GPCRs (Fig. 1c). This suggests that the CFEM domain is not inherently essential for the functionality of Pth11-like GPCRs in fungi.

**Table 1:**
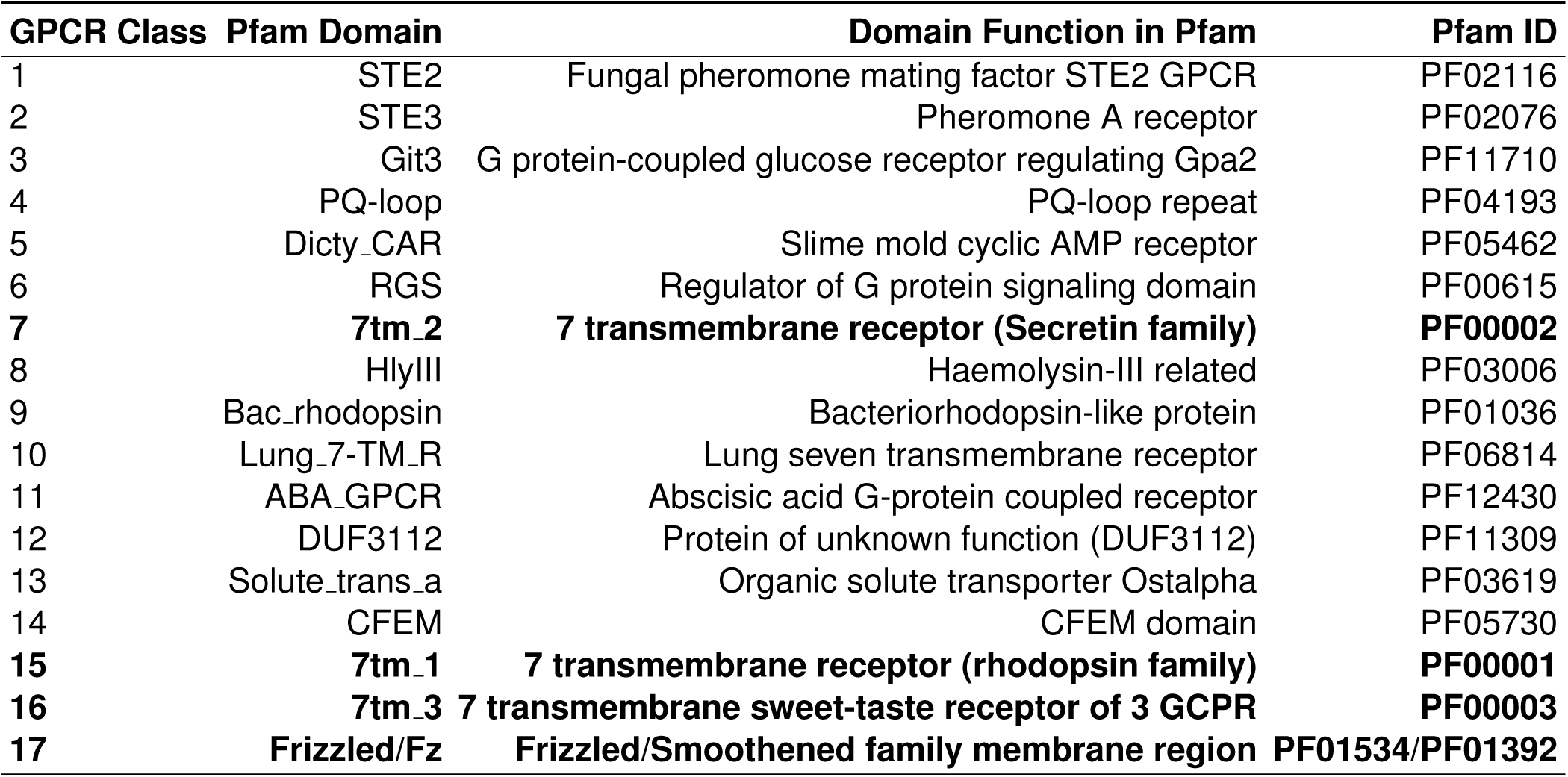
List of all GPCR classes, related Pfam domains, and their functions. The four mammalian GPCR classes detected in fungi are highlighted in bold.

### Identification of mammalian GPCR homologs in fungal species

To identify mammalian GPCR homologs in fungi, we searched for functional Pfam domains corresponding to five human GPCR classes (Rhodopsin, Glutamate, Frizzled, Taste2, and Adhesion) across 1,357 fungal species. After filtering for sequences containing seven TMHs, we identified 2,089 mammalian GPCR homologs in 594 fungal species, distributed across three mammalian GPCR families: Rhodopsin, Glutamate, and Frizzled, which we suggest to add to fungal GPCR classification as classes 15,16, and 17, respectively (Fig. 1c-d).

Unlike the 14 fungal GPCR classes, which are predominantly enriched in Ascomycota, mammalian GPCR homologs are primarily found in EDF. EDF species have a higher average number of mammalian homologs per species and display a substantially fewer missing or decreased GPCRs at both the species and GPCR class levels, compared to Ascomycota and Basidiomycota (Fig. 1b). The majority of mammalian homologs were found in the Rhodopsin class (Class 15), with 984 sequences (47.1%), followed by 918 sequences (43.9%) in the Glutamate class (Class 16), with the remaining 187 (8.9%) in the Frizzled class (Class 17) (Fig. 1d). Nearly no Glutamate and Frizzled homologs were found in Ascomycota and Basidiomycota. Large number of Rhodopsin homologs were found in Entomophthoromycotina, a subphylum containing arthropod pathogens and soil-and litter-borne saprobes. Glutamate homologs are enriched in Blastocladiomycota and Chytridiomycota, phyla of fungi characterized by their production of zoospores with a single posterior flagellum often found in aquatic habitats, acting as parasites of plants. In summary, we identified mammalian GPCR homologs from four distinct classes, including the Secretin class, which shares the 7tm 2 domain with Class 7 in fungal GPCRs. Interestingly, the Adhesion and Taste2 classes appear to be absent in fungi. This demonstrates some evolutionary conservation of key GPCR families between mammals and fungi, especially in EDF.

### Comparative analysis of Pth11-like GPCRs

The Pezizomycotina subphylum, comprising 650 species with sequenced genomes, exhibits a significant prevalence of Pth11-like GPCRs (Class 14) relative to other GPCR classes, accounting for two-thirds of all fungal GPCRs. Nonetheless, these GPCRs, found in seven fungal groups within Pezizomycotina, display variable abundance (Fig. 2), up to tenfold. In order to examine the evolutionary relationships of Pth11-like GPCRs among closely related Pezizomycotina species that exhibit significant variations in receptor abundance, a comparative phylogenetic analysis of 12 selected representative species of Sordariomycetes (Table 2) was conducted.

**Figure 2:**
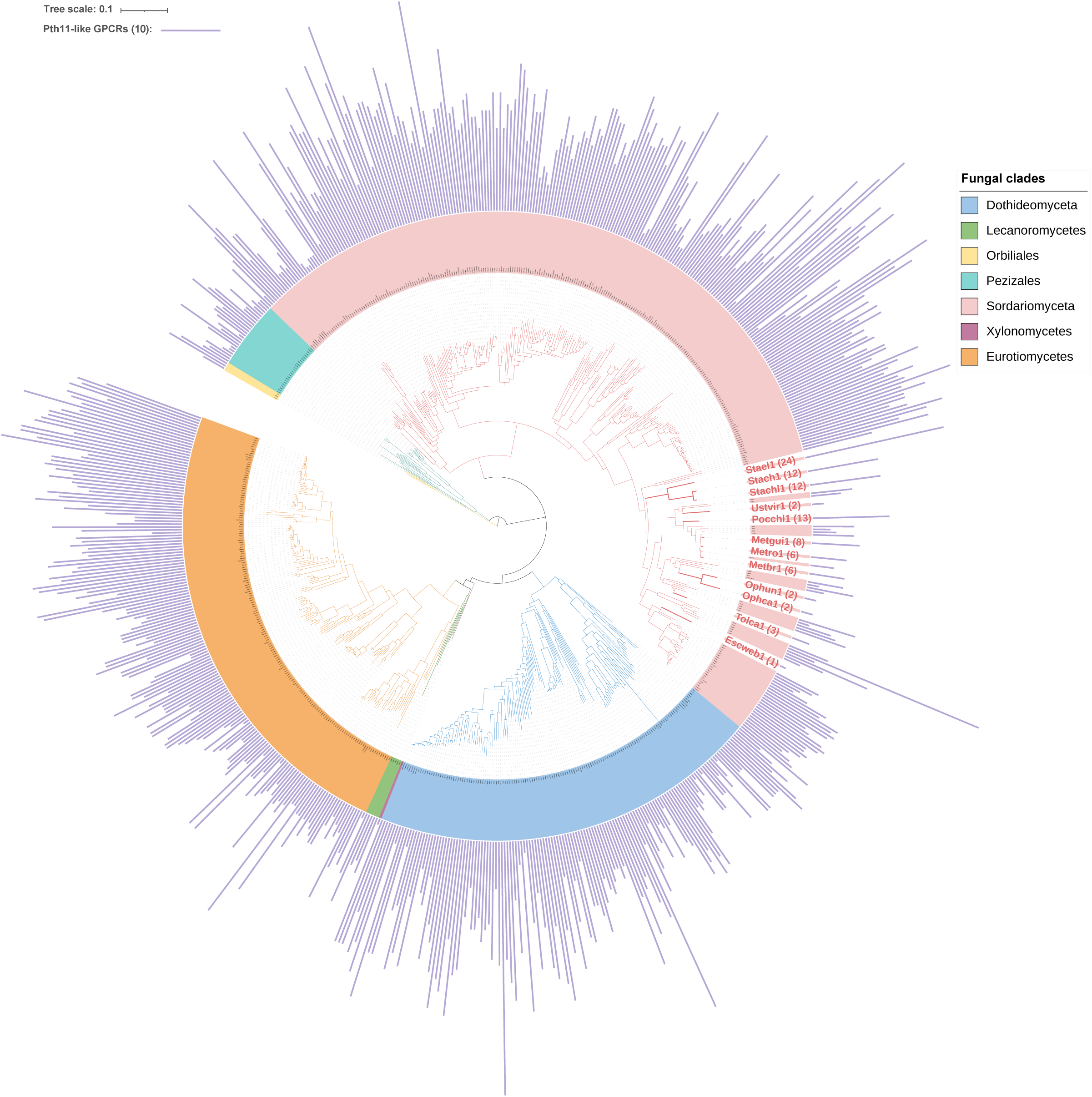
Phylogenetic tree of Pth11-like GPCRs across 530 species from five fungal classes and two orders (Orbiliales and Pezizales) within Pezizomycotina. Species from different classes are color-coded according to the legend. Twelve species from Sordariomyceta are highlighted in bold with enlarged font (Table 2), selected for their phylogenetic proximity and notable variation in Pth11-like GPCR abundance. The number in parentheses indicates the abundance of Pth11-like GPCRs. The purple bar on the outside of the circle represents the abundance of Pth11-like GPCR in each species, varying from 1 to 43 in individual genomes.

**Table 2:** Pth11-like GPCR characteristics in twelve selected species in Pezizomycotina. Twelve selected species are arranged in ascending order of Pth11-like GPCR abundance. The number of Pth11-like GPCR duplication and loss events, identified using Notung[34], along with the number of expanded and contracted gene families in their genomes, as determined by CAFE[35], are displayed.

| Species | MycoCosm portal ID | Pth11-like GPCR Abundance | Gene Duplications | Gene Losses | Expanded Gene Families | Contracted Gene Families |
| --- | --- | --- | --- | --- | --- | --- |
| *Escovopsis weberi* | Escweb1 | 1 | 0 | 14 | 115 | 2816 |
| *Ophiocordyceps camponoti-rufipedis* | Ophca1 | 2 | 0 | 1 | 113 | 617 |
| *Ophiocordyceps kimflemingae* | Ophun1 | 2 | 0 | 1 | 162 | 175 |
| *Ustilaginoidea virens* | Ustvir1 | 2 | 0 | 11 | 99 | 3361 |
| *Tolypocladium capitatum* | Tolca1 | 3 | 0 | 2 | 222 | 2400 |
| *Metarhizium brunneum* | Metbr1 | 6 | 0 | 4 | 89 | 451 |
| *Metarhizium robertsii* | Metro1 | 6 | 0 | 4 | 179 | 138 |
| *Metarhizium guizhouense* | Metgui1 | 8 | 0 | 4 | 265 | 433 |
| *Stachybotrys chartarum* | Stach1 | 12 | 0 | 3 | 343 | 484 |
| *Stachybotrys chlorohalonata* | Stachl1 | 12 | 0 | 3 | 136 | 560 |
| *Pochonia chlamydosporia* | Pocchl1 | 13 | 1 | 2 | 869 | 742 |
| *Stachybotrys elegans* | Stael1 | 24 | 4 | 7 | 1074 | 539 |

Initially, we examined the duplication and loss of Pth11-like GPCRs among these 12 species utilizing Notung [34], uncovering 42 duplication and 108 loss events. *Escovopsis weberi* demonstrated the highest losses of Pth11-like GPCRs (Table 2). Species with a high abundance of GPCRs typically experienced fewer loss events and more duplications. This suggests that the variation in Pth11-like GPCR abundance is driven by losses with some contributions from duplication events. This also indicates that the ancestor for these species already had high number of Pth11-like GPCRs. We further analyzed genome expansion and contraction across these species using CAFE [35]. *E. weberi*, *Tolypocladium capitatum*, and *Ustilaginoidea virens* demonstrated a significant quantity of contracted gene families, accompanied by a comparatively lower number of expanded families. Conversely, species exhibiting a significant abundance of Pth11-like GPCRs, including *Pochonia chlamydosporia* and *Stachybotrys elegans*, demonstrated a comparatively higher number of expanded gene families. This suggests a correlation between the expansion or contraction of Pth11-like GPCRs and the overall dynamics of the genome. The genome size contraction of species with a low abundance of Pth11-like GPCRs may be linked to alterations in their lifestyle.

Gene Ontology (GO) analysis was performed on all genes, excluding GPCRs, across the six species that exhibit a Pth11-like GPCR abundance difference of no less than six-fold. In *E. weberi*, significantly expanded genes (*P* < 0.05) were enriched in functions related to gene expression regulation and enzymatic activity modulation (Fig. 3a). On the other hand, contracted genes were predominantly linked to transmembrane transport and hydrolase activity. Comparable enrichment patterns were noted in *T. capitatum* and *U. virens* (Supplementary file 2). In contrast, *S. elegans* displayed expansion in genes involved in transmembrane transport and oxidoreductase catalytic activity, with no significant contraction in gene sets (Fig. 3b). Comparable results were noted for *P. chlamydosporia* and *Stachybotrys chartarum*.

**Figure 3:**
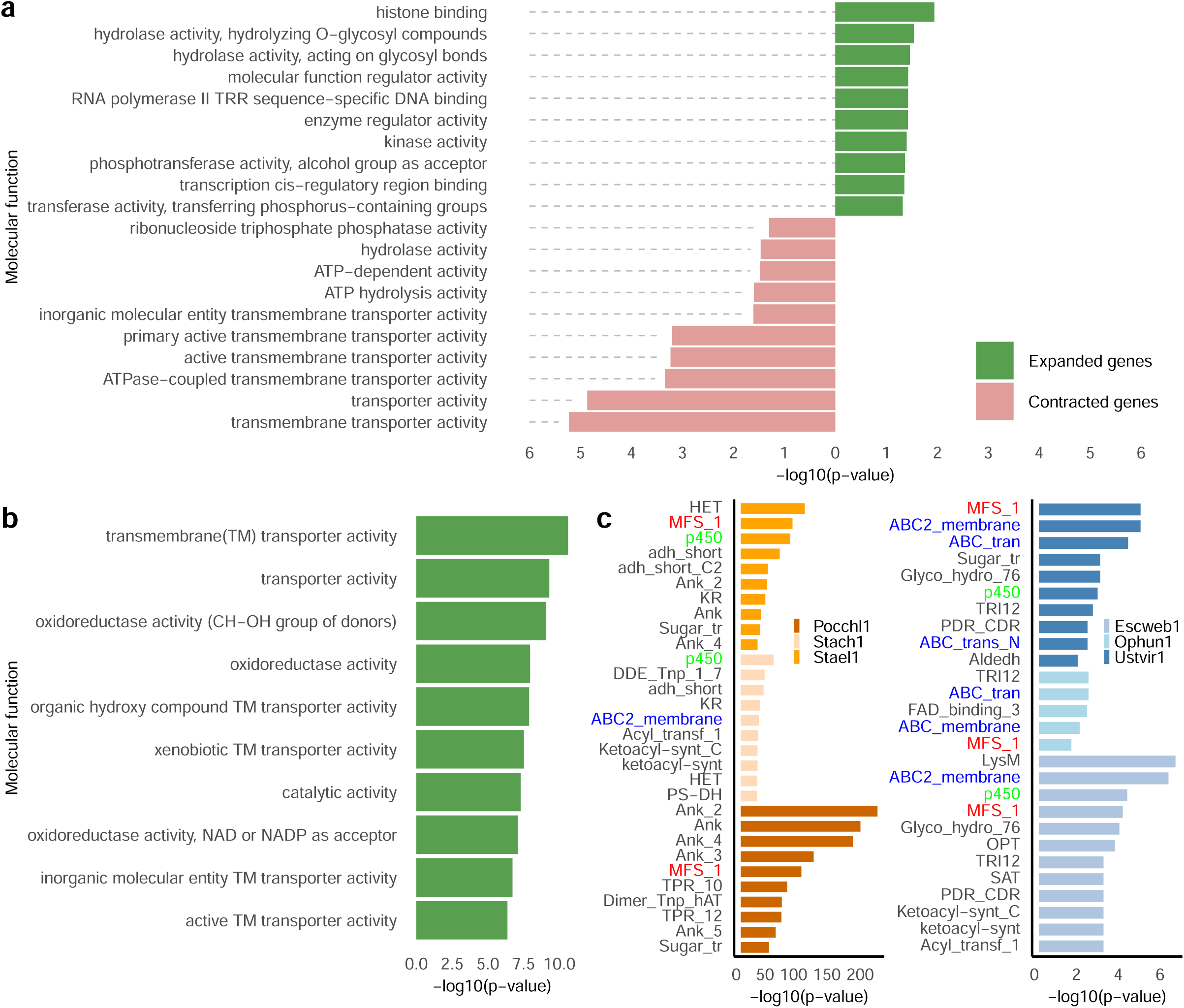
Functional enrichment and Pfam domain analysis across fungal species based on contracted and expanded gene families identified using CAFE. **a** Top ten molecular functions for both significantly expanded and contracted genes in the GO enrichment analysis for *Escovopsis weberi*. **b** Top ten molecular functions for significantly expanded genes in GO enrichment analysis for *Stachybotrys elegans*. No significantly contracted genes were identified in *Stachybotrys elegans*. **c** Pfam domain enrichment analysis: The left panel shows the top significantly expanded Pfam domains in *Pochonia chlamydosporia* (Pocchl1), *Stachybotrys chartarum* (Stach1), and *Stachybotrys elegans* (Stael1). The right panel shows the top significantly contracted Pfam domains in *Escovopsis weberi* (Escweb1), *Ophiocordyceps kimflemingae* (Ophun1), and *Ustilaginoidea virens* (Ustvir1). Common Pfam domains across species are bolded. A two-sided t-test was performed to identify significant expanded or contracted genes, with p <0.05 considered significant. The Benjamini-Hochberg method was used to adjust p-values for multiple testing.

Additionally, Pfam domain enrichment analysis indicated that the contracted genes were predominantly enriched in the MFS 1 domain in *E. weberi*, *T. capitatum*, and *U. virens* (Fig. 3c). The MFS (major facilitator superfamily) comprises a large group of plasma membrane proteins that function as transmembrane transporters for a wide range of substances, and these transporters are both abundant and diverse in fungi [36]. These three species also exhibited gene contraction in domains related to ABC (ATP-binding cassette) transporters, which facilitate the transport of natural metabolites and xenobiotics—such as antifungal compounds—through ATP hydrolysis, playing a key role in antifungal resistance [37]. Moreover, *E. weberi* and *U. virens* demonstrated a marked reduction in cytochrome P450 enzymes. Conversely, *S. chartarum* and *S. elegans* demonstrate considerable expansion in genes linked to the MFS 1 domain, but both *S. elegans* and *P. chlamydosporia* indicate an enrichment of genes pertaining to cytochrome P450. The expanded gene sets in these species are also enriched in Pfam domains such as HET, adh short, and Ank, which are involved in cellular processes and metabolic regulation.

Furthermore, given that Pth11-like GPCRs, which detect cellulose and plant cell wall components, trigger responses promoting fungal infection [23, 38], we investigated their association with key proteins involved in secondary metabolism across multiple species.In a total of 212 species, Pth11-like GPCRs are positioned adjacent to biosynthetic gene clusters that participate in secondary metabolism. These GPCRs are located most frequently in close proximity to polyketide synthase (PKS) and nonribosomal peptide synthetase-like (NRPS-like) genes, which implies that they may play a role in the regulation of secondary metabolite production (Supplementary file 1 Table S3).

### Novel GPCRs predicted using machine learning

After excluding sequences previously identified as putative GPCRs by HMM-based searches, 141,666 protein sequences, containing seven TMHs, remain unidentified as GPCRs among 1,355 fungal species. MCL clustering of these sequences resulted in 1,969 clusters, with an average size of 71.5 sequences, and the biggest cluster of 4,046 sequences. Pfam domain identification was conducted on the ten largest clusters to elucidate the functional roles of the sequences. The analysis indicated that the majority of sequences within specific clusters have identical Pfam domains, whilst other clusters predominantly lacked identifiable Pfam domains, indicating proteins of unknown functions (Fig. 4a). Given that 96.22% of the sequences within the largest cluster contained the DUF6534 domain, we suggest that these sequences might exhibit GPCR functionality, potentially representing a new class of GPCRs.

**Figure 4:**
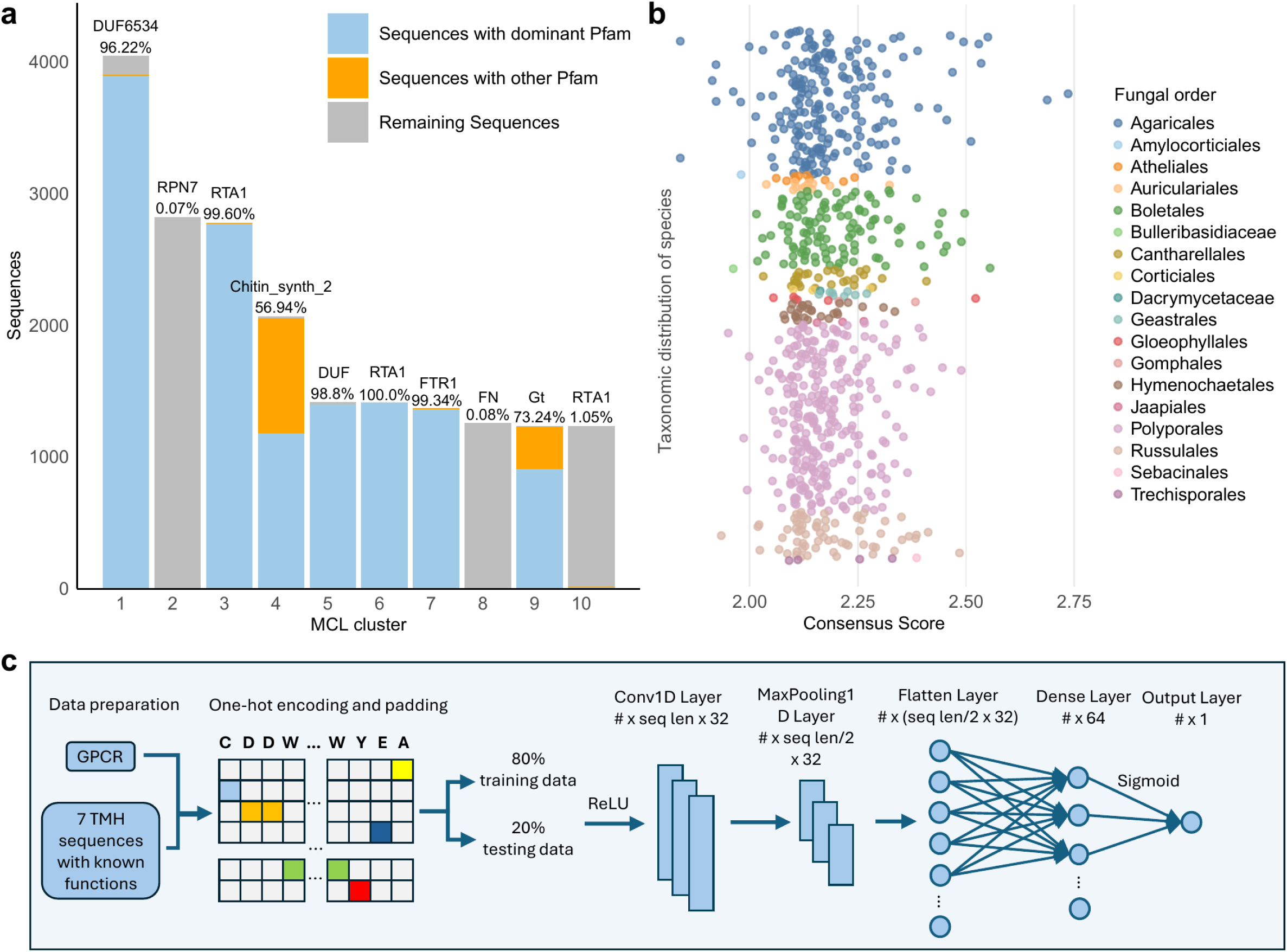
Identification of novel GPCRs. **a** Top ten largest MCL clusters for sequences with seven TMHs, excluding the 31,493 newly identified GPCRs. The blue portion of each bar represents the number of sequences with the dominant Pfam domain in that cluster, the orange portion represents sequences with other Pfam domains, and the grey portion represents the remaining sequences. The dominant Pfam domain and its percentage are displayed on top of each bar. FN in cluster 8 refers to Flavi NS4A and Gt in cluster 9 refers to Glyco trans 2 3. **b** Consensus score distribution (sum of self-built CNN, ProteInfer, and DeepFRI models) across 222 species in 18 fungal orders. **c** Architecture of the self-built 1D-CNN model. Newly identified 16,055 GPCR sequences with GPCR-related Pfam domains are used as the positive dataset, while 17,997 sequences with seven TMHs in MCL clusters dominated by Pfam domains unrelated to GPCR functions are used as the negative dataset. Sequences are transformed into one-hot encoding and padded. 80% of sequences are used as training data, and 20% as testing data. The sequences are input into a convolutional 1D layer with 32 filters, a kernel size of 3, and ReLU activation. This is followed by a max-pooling layer with a pool size of 2. The flattened output is passed through a fully connected layer with 64 neurons (ReLU activation), and finally, a sigmoid-activated output layer is used for binary classification.

To test this hypothesis, machine learning algorithms ProteInfer and DeepFRI were employed to classify whether the 3893 sequences containing the DUF6534 domain in the largest MCL group are potential novel GPCRs. ProteInfer predicted 1793 sequences with GPCR activity with a confidence score exceeding 0.8 and within them, 1637 sequences were predicted to belong to the GPCR A Pfam clan, which includes numerous members of the rhodopsin superfamily. Whereas DeepFRI identified 2204 sequences that are predicted to have GPCR activity and only 25 sequences of them that have GPCR activity with a confidence score above 0.5 (considered high confidence), there exist sequences with DeepFRI scores <0.2 that represent correct predictions [39]. The substantial divergence between the outcomes of ProteInfer and DeepFRI indicates a need for a novel classifier to assess the GPCR activity of proteins containing the DUF6534 domain.

Consequently, we designed a one-dimensional convolutional neural network (CNN) model designed to detect putative new GPCRs among proteins anticipated to possess 7-9 TMHs (Fig. 4c). We evaluated 8 or 9 TMHs in addtion to 7 TMHs proteins because Class 11 GPCRs possess 9 TMHs [22], while also considering possible discrepancies in the TMHMM prediction. The model was trained on a dataset that included 17,996 sequences from 14 GPCR classes, as well as a negative dataset of 18,221 sequences that featured 7 TMHs but were linked to non-GPCR functions (see Methods for details). An F1 value of 0.97 was obtained during the model validation process, indicating exceptional performance in the classification of GPCRs and non-GPCRs. Utilizing this model, we predicted 71.3% of sequences within the largest clusters as GPCRs. Subsequently, we integrated confidence scores from ProteInfer (scores >0.8), DeepFRI (scores >0.2), and the CNN model (scores >0.9) to derive a consensus prediction for GPCR activity. As a result, consensus GPCR prediction was obtained for 767 sequences from 222 fungal species, all of which are in the Agaricomycotina subphylum in Basidiomycota (Fig. 4b). These species included *Dioszegia hungarica* from Tremellomycetes and *Calocera cornea* from Dacrymycetes and the rest from Agaricomycetes. The orders Polyporales, Agaricales, and Boletales contain the most sequences. Notably, Agaricales has the highest number of sequences with scores above 2.50. Since most consensus scores range between 2 and 2.50, establishing a threshold for accurate prediction of a novel GPCR is challenging. Therefore, AlphaFold Multimer [33] was used to evaluate the interaction potential between the predicted GPCRs and the G-alpha proteins to *in silico* validate their function.

### Predicted interactions between potential novel GPCRs and G-alpha proteins

After identifying the most promising novel GPCRs across 222 species, the G-alpha proteins were quantified for each species, showing considerable variation in their quantities. However, there is no clear relationship between the quantity of G-alpha proteins and the total number of GPCRs (Pearson correlation=-0.256). To identify interactions between novel GPCRs and G-alpha proteins using AlphaFold Multimer [33], we conducted preliminary tests with known functional pairs and mismatched counterparts. There are two well-characterized GPCR-G-alpha protein pairs involved in distinct pathways controlling mating in *S. cerevisiae*. The first pair, related to nutrient availability, involves Gpr1 and Gpa2 [12]. Gpr1 functions as a carbon receptor sensing glucose, causing a conformational change that activates the G*α*-protein Gpa2. The second pair involves Ste2 interacting with Gpa1 in the pheromone sensing pathway [40]. Leveraging published research, we note that Gpa1 and Gpa2 are the only two G*α* subunit genes in the *S. cerevisiae* genome responsible for distinct pathways [11]. Additionally, 17 well-characterized GPCR-G-alpha protein pairs in two other fungal species (*C. albicans*, and *Cryptococcus neoformans*), one plant species (*Arabidopsis thaliana*) and humans were included as positive control (Supplementary file 1 Table S4). Since Gpa1 can productively couple with the *α*-factor receptor Ste2, whereas Gpa2 cannot [41], we exchanged the G*α* proteins between the GPCRs to create two pairs as negative controls, along with 23 extra pairs composed of non-GPCR proteins (with unrelated Pfam domains) from *S. cerevisiae* paired with Gpa1 or Gpa2 as negative controls. AlphaFold Multimer was conducted for these 44 protein pairs. The ACM2-GNAI2 pair in humans achieved the highest confidence score of 0.77 (0.8 ipTM + 0.2 pTM) [33]. In fungi, the highest score was 0.70 for the Ste2-Cag1 pair from *Candida albicans*. In contrast, the best score among the non-interacting pairs was 0.52. On average, interacting pairs had a mean confidence score of 0.62, significantly higher than the mean score of 0.32 for non-interacting pairs (Fig. 5a). A two-sided t-test revealed that AlphaFold Multimer scores are significantly different between interacting and non-interacting complexes (p = 2.44e-14). This demonstrates AlphaFold Multimer’s strong performance in distinguishing between interacting and non-interacting complexes.

**Figure 5:**
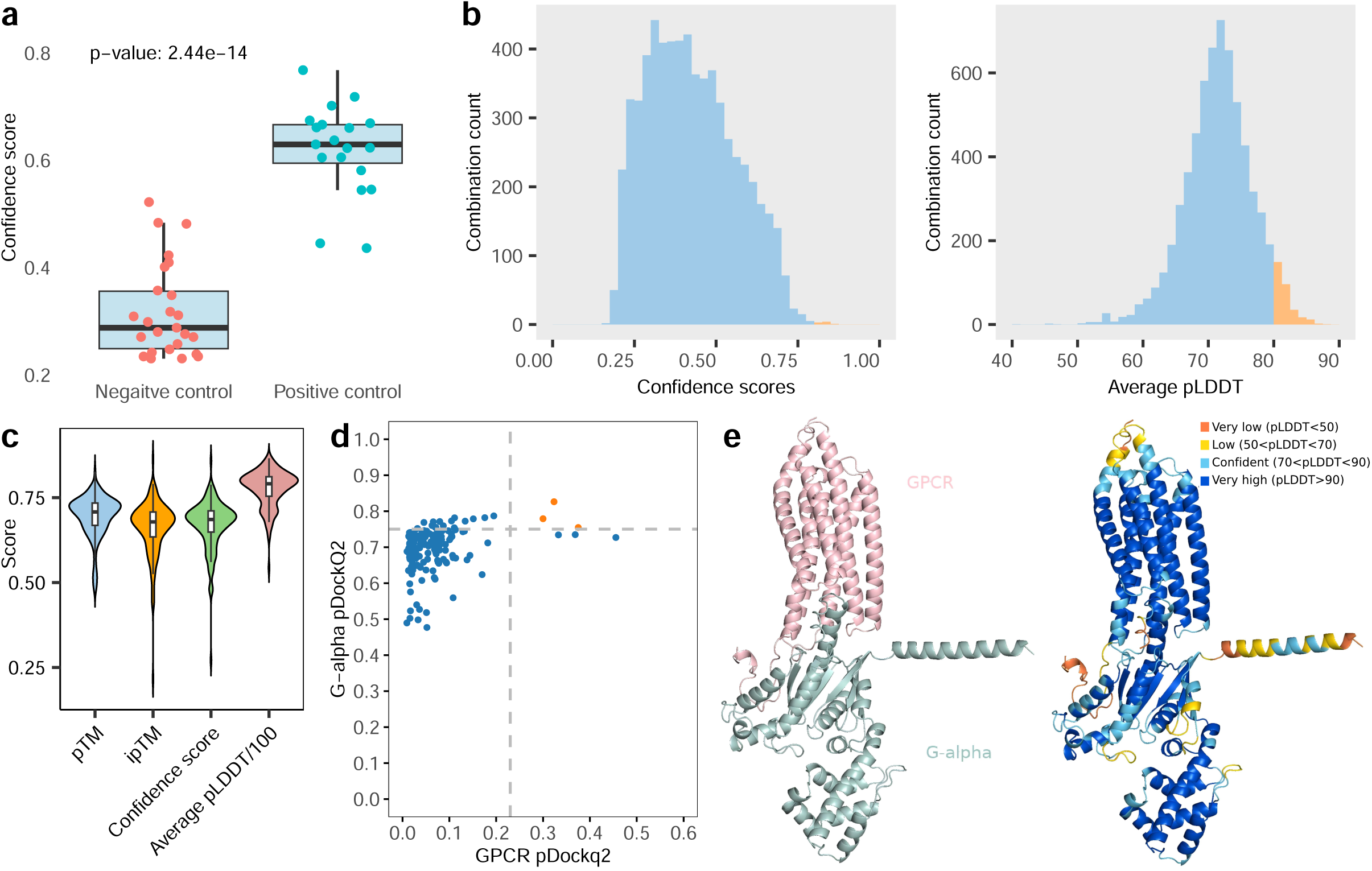
Evaluation of GPCR-G-alpha interactions with AlphaFold Multimer. **a** Comparison of AlphaFold Multimer confidence scores between interacting and non-interacting protein pairs [33]. For each combination, five AlphaFold Multimer models were generated and the highest scores are presented. The median, 25th, and 75th percentiles are shown. **b** Distribution of confidence scores (calculated as 0.8 × iPTM + 0.2 × PTM) and the average pLDDT score across all GPCR and G-alpha combinations. Combinations with a score higher than 75 are highlighted in orange. Similarly, combinations with an average pLDDT score above 80 are considered high-confidence and are also highlighted in orange. **c** Evaluation metrics for the top pair from 155 species, based on four AlphaFold Multimer results. **d** pDockQ2 scores for both GPCR and G-alpha are shown. Interactions with a minimum pDockQ2 score greater than 0.23 are considered high-confidence. **e** The structure of the GPCR and G-alpha interaction with the highest confidence, as predicted in *Galerina marginata*. The corresponding pLDDT scores of the structure are also displayed.

However, the scores for some interacting complexes are moderate for known interactions but lack detailed information, as they do not include per-residue local confidence (pLDDT) [42] provided by AlphaFold 2 or the global confidence measure provided by the predicted aligned error (PAE) in AlphaFold-Multimer. Therefore, we implemented the pDockQ2 score [43], which is capable of assessing AlphaFold Multimer results that incorporate per-residue and PAE information. pDockQ2 scores of 0.262 and 0.394, respectively, were obtained for Gpr1 and Gpa2, with scores exceeding 0.23 suggesting high-confidence interactions. The scores of Ste2 and Gpa1 in the second pair were 0.307 and 0.409, respectively. The effectiveness of the pDockQ2 score in identifying protein interactions was underscored by the unambiguous distinction of the pDockQ2 scores for the non-interacting pairs, with 0.022 for Gpr1-Gpa1 and 0.021 for Ste2-Gpa2.

Then, we employed AlphaFold Multimer to assess all 6,376 possible pairings of new GPCRs and G-alpha proteins across 155 fungal species. By choosing the maximum confidence level for each combination, we determined that the overall average confidence score is 0.425. Significantly, merely nine combinations possess confidence levels above 0.8, so categorizing them as high-confidence predictions. All nine derive from *Galerina marginata*, a white-rot fungus belonging to the Cortinariaceae family. The mean pLDDT for all combinations is 71.5, with 312 combinations exhibiting pLDDT values over 80 (Fig. 5b), suggesting possible interactions between the new GPCRs and G-alpha proteins.

There was considerable variation in the highest confidence scores for different combinations within a single species (Supplementary file 1 Table S5). We selected the combination with the highest confidence score for each species and summarized their evaluation metrics. The pTM and ipTM have similar average scores with ipTM having larger variance (Fig. 5c). Majority of the average pLDDT concentrated at 80, which can be considered as high confidence. Then we calculated the pDockQ2 score for the species. Six species exhibited a minimum pDockQ2 > 0.23, and three of these had a pTM > 0.75 (Fig. 5d), indicating high-confidence combinations. The three species identified in this study are *G. marginata* (Fig. 5e), *Bjerkandera adusta*, a known plant pathogen that causes white rot in both live trees and dead wood, and *Coprinopsis sp. MPI-PUGE-AT-0042 v1.0*, which colonizes the roots of healthy *A. thaliana* plants grown in natural soil after surface sterilization. All three species belong to the Agaricomycetes class.

## Discussion

This study offers an in-depth investigation of GPCR diversity among more than 1,300 fungal species, utilizing established fungal GPCR classification consisting of 14 known fungal classes and additional three mammalian classes not previously detected in fungi, uncovering notable variation in GPCR abundance and class distribution among Ascomycota, Basidiomycota, and EDF. The fungal groups Olpidiomycotina, Apansporoblastina, and Pansporoblastina which encompass parasitic fungi in EDF, were entirely devoid of 17 recognized GPCR categories. The findings indicate that specific early-diverging fungal lineages may have bypassed the evolutionary development of GPCR-mediated signaling pathways. The lack of GPCRs in these populations may suggest an evolutionary adaptation in which alternative membrane proteins or other mechanisms perform similar signaling roles. This may indicate a diminished dependence on GPCRs as a result of ecological specialization or variations in membrane structure, emphasizing the divergent evolutionary pressures that are exerting themselves on these parasitic fungi. Ascomycota exhibited the highest GPCR abundance and diversity in comparison to Basidiomycota and EDF, particularly with the dominance of Pth11-like GPCRs, while mammalian classes Rhodopsin and Glutamate (fungal newly added classes 15 and 16) dominated in EDF.

At the GPCR class level, Class 1 GPCRs, represented by STE2-like pheromone receptors, were found to be exclusive to Ascomycota, suggesting that this class evolved after the divergence of Ascomycota from EDF. Ascomycota either developed these receptors independently or expanded an ancestral signaling pathway into the Class 1 GPCRs. The absence of Class 1 GPCRs in Basidiomycota may suggest that the pheromone signaling pathways involving Class 1 GPCRs are either absent or have been supplanted by alternative mechanisms. This is further illustrated by the predominance of Class 2 GPCRs in both Ascomycota and Basidiomycota. The evolutionary proliferation of Class 2 GPCRs in Basidiomycota, particularly within Agaricomycetes, indicates that these fungi may exhibit adaptations to complex mating systems, presumably influenced by selective forces aimed at improving partner identification and reproductive success across many ecological niches. Additionally, the absence of GPCR classes 5, 6, and 7 in Basidiomycota, in conjunction with their distinctive occurrence in Ascomycota and EDF, suggests that evolutionary forces are specific to a particular phylum. These receptors may have been essential for specific environmental detection or developmental processes that were unique to Ascomycota and EDF. Their absence in Basidiomycota indicates a possible loss attributable to functional redundancy, wherein other signaling pathways may have developed to perform analogous functions.

The functional relevance of the 14 classes of identified GPCRs is confirmed by presence of GPCR-related Pfam domains in over 98.9% of sequences with Pfam domains. Interestingly, the CFEM domain, which is normally linked with pathogenicity, was found in a small subset of class 14 GPCRs, implying that it is not required for Pth11-like GPCR function. Pth11-like GPCRs make for almost one-third of all discovered GPCRs, and they are all found exclusively in Pezizomycotina. This is consistent with prior findings demonstrating that Pth11-like GPCRs are exclusive to Pezizomycotina [27] but is confirmed here for much larger number of genomes.

Furthermore, we found 2,089 mammalian GPCR homologs among 594 fungal species, thereby augmenting the known fungal GPCR repertoire by three additional classes beyond the previously recognized 14 classes. The newly found classes, analogous to mammalian Rhodopsin, Glutamate, and Frizzled GPCRs, constitute the inaugural reported increase of fungal GPCR taxonomy to a total of 17 unique classes. The distribution of these mammalian GPCR homologs differs from the pattern shown in the 14 previously identified fungal classes, which are mainly located in Ascomycota. Mammalian GPCR homologs are predominantly concentrated in EDF lineages. Rhodopsin homologs are markedly concentrated in Entomophthoromycotina, whilst Glutamate homologs are prominently abundant in both Blastocladiomycota and Chytridiomycota. This pattern indicates a compelling evolutionary connection between EDF species and mammals, possibly signifying a common ancestral GPCR repertoire. The presence of these homologs implies that EDF lineages may have preserved ancient GPCRs that were maintained as a result of selective pressures associated with parasitism or specific ecological niches. At the same time, fungi have evolved a multitude of lineage-specific GPCRs over the course of evolution, which is likely due to their adaptations to a variety of environmental conditions. These led to significant reductions of the previously recognized 14 GPCR classes in EDF species, whereas they are prominently represented in Ascomycota and Basidiomycota. These discoveries underscore the evolutionary adaptability of the GPCRs within fungal lineages and emphasize the importance of ecological and evolutionary forces in influencing the diversity of signaling systems in fungi.

Our analysis further indicates considerable diversity in the prevalence of Pth11-like GPCRs among over 600 species within the Pezizomycotina subphylum. Utilizing comparative phylogenetic analysis, gene duplication and loss event detection, with genome expansion and contraction investigations, we aim to elucidate the evolutionary mechanisms responsible for the variation in Pth11-like GPCR abundance. Our findings suggest that variations in the abundance of these receptors are closely associated with gene duplication and loss events, along with larger genomic dynamics, including the expansion and contraction of gene families. Species possessing a greater quantity of Pth11-like GPCRs, such as *P. chlamydosporia* and *S. elegans*, demonstrated an increased number of gene family expansions, suggesting a possible correlation between receptor prevalence and genomic dynamics. Conversely, species with a diminished number of Pth11-type GPCRs, such as E. weberi, exhibited a greater degree of gene family contraction.

Moreover, by comparing these findings with functional annotations, including Gene Ontology (GO) and Pfam domain enrichment, we examined the potential relationship of Pth11-like GPCRs with secondary metabolism and host-pathogen interactions. The proliferation of Pfam domains like MFS 1 and cytochrome p450 in species with elevated Pth11-like GPCR counts indicates improved abilities for environmental adaptation, nutrient acquisition, and stress response, whereas species with diminished GPCR abundance may have developed more efficient and specialized interactions with their hosts.This divergence presumably corresponds to markedly different evolutionary pressures, with parasitic species such as *S. elegans* evolving more complex regulatory and signaling systems in association with a parasitic lifestyle, while *E. weberi* may have adapted to a more specialized ecological niche which still relies on its host for metabolic requirements. Pth11-like GPCRs may thereby affect the intricacy of pathogenicity. Certain investigations indicate an association between elevated Pth11-like GPCR quantities and an expanded host range [44]. However, the case of *S. elegans*—which shows the highest Pth11-like GPCR abundance yet exhibits narrow host specificity to *R. solani AG-3*—indicates that Pth11-like GPCR abundance may be more closely associated with specialized parasitic strategies and pathogenic intricacy. The parasitic lifestyle of *S. elegans*, which involves the production of cell wall-degrading enzymes, intracellular colonization, and the expression of pathogenicity-associated genes, is likely supported by the expansion of signaling-related genes [45]. In contrast, *E. weberi*, a specialized parasite of the fungal cultivar *Leucoagaricus gongylophorus*, exhibits low Pth11-like GPCR abundance and appears to depend heavily on its host for signaling and metabolic processes. This reliance may have led to its genome reduction [46] and a more simplified host interaction mode. Such dependency illustrates an evolutionary trade-off, where the parasite has streamlined its genome to adapt to its highly specialized ecological niche within ant colonies. Moreover, 212 species have Pth11-like GPCRs located near biosynthetic gene clusters, particularly adjacent to PKS and NRPS-like genes, implying a role in regulating secondary metabolism.

Additionally, we identified a significant number of proteins consisting of seven TMHs that may represent novel GPCRs while assessing the established GPCR distributions among fungal species. These can be predicted using a variety of protein function prediction techniques. Our research utilized machine learning and clustering methodologies to analyze a significant collection of unclassified protein sequences that included seven TMHs. Consequently, we were able to predict the emergence of new GPCRs in 222 fungal species. The identification of 767 sequences with consensus predictions of GPCR activity was facilitated by the integration of multiple machine learning methods, such as ProteInfer, DeepFRI, and a custom CNN model. The clustering and domain analysis reveal that a considerable proportion of these sequences belong to an uncharacterized GPCR 14 family, especially those featuring the DUF6534 domain. These novel GPCRs were predominantly detected in species within the Basidiomycota and Agaricomycotina, namely in the orders Polyporales, Agaricales, and Boletales. Discrepancies in protein function prediction results between ProteInfer and DeepFRI highlight the need for further refinement of GPCR classification models. Although the majority of consensus scores fell within a narrow range, making it difficult to establish a definitive threshold for high-potential GPCRs, our findings suggest that these novel GPCR candidates may represent a previously unrecognized class of signaling proteins. In order to evaluate their potential for interaction with G-alpha proteins, which is essential for determining their biological function, additional structural validation was performed using AlphaFold Multimer. In the subsequent investigation into the potential interactions between these novel GPCRs and G-alpha proteins, AlphaFold Multimer was used to predict 3D interactions across 155 fungal species. In our analysis of 19 well-characterized GPCR-G-alpha protein pairs in a variety of species, we found that the efficacy of incorporating both AlphaFold Multimer and pDockQ2 scores in distinguishing between interacting and non-interacting complexes was demonstrated, thereby bolstering our confidence in our approach. Out of the 6,376 combinations that were assessed, nine high-confidence interactions were identified, all of which originated from *G. marginata*. This indicates that these novel GPCRs may have a potential function in fungal signaling pathways. The variation in interaction confidence across species, with six species exhibiting pDockQ2 scores above 0.23 and three species displaying both high pTM and pDockQ2 scores, points to the specificity of GPCR-G-alpha protein interactions in different fungal contexts. It is important to note that the high-confidence interactions predicted in *G. marginata*, *B. adusta*, and *Coprinopsis sp.* indicate that these GPCRs may be essential for plant-fungal interactions, particularly in the processes of wood decomposition and root colonization. The high pLDDT scores and structural models generated by AlphaFold Multimer establish a foundation for the future experimental validation and functional characterization of these novel GPCRs in fungal signaling and pathogenesis.

Our analysis of more than 1,300 fungal genomes highlights the extraordinary diversity of GPCRs across fungal tree of life. We have expanded the fungal GPCR classification by adding three classes previously identified only in mammals and predicting novel GPCR-related domains. In addition, we discovered that Pth11-like GPCR abundance varies significantly among species in Pezizomycotina, primarily due to gene losses during evolution. Our work sets a stage for the functional validation of GPCRs and further inquiry into their biological functions.

## Materials and Methods

### Source of the data

The query sequences of all 14 classes of GPCRs were sourced from previous literature [47]. The genomes of 1,357 fungi species are publicly available in MycoCosm [48]. The genome scaffolds for *Carpinus fangiana*, *Quercus suber*, *Cichorium endivia* and *Chlamydia trachomatis* were obtained from NCBI (https://www.ncbi.nlm.nih.gov/genome/). Phylogenetic tree of the Pezizomycota was obtained from the MycoCosm portal [48].

### Identification of putative GPCRs across fungal species using known fungal GPCR classification

In this comprehensive analysis, 67 GPCR query sequences, spanning all 14 recognized classes, were subjected to BLASTP 2.8.1+ [21] searches against genomes of 1,355 species from the MycoCosm database [48]. Searches were constrained to an E-value ˂1E-05 and required hits to cover >75% of both the query and MycoCosm protein entries. The number of TMHs of the candidate GPCRs were verified using TMHMM [49], retaining only sequences with 6-7 TMHs, to account for TMHMM’s limitations in accurately determining helix counts. For Class 9 GPCRs, we maintained the sequences with 7 to 9 TMHs because Class 9 GPCRs were originally found with 9 TMHs [22]. Multiple sequence alignments were executed with MAFFT version 6.717 [50]. Then regions lacking conservation were excised using trimAl [51]. Hidden Markov Models (HMMs) were constructed for 14 GPCR classes using ‘hmmbuild’ from HMMER 3.2 [52], and ‘hmmsearch’ was employed against publicly available proteomes from MycoCosm databass to identify potential homologs and results were filtered with an E-value threshold of <1e-05. A second round of TMHMM analysis was applied to the sequences obtained from ‘hmmsearch’ to filter out those not meeting the TMHs criteria. Finally, Pfam domain identification isolated sequences containing fungal GPCR-functional domains (Table 1), thereby delineating a subset of putative GPCRs within a diverse array of fungal species.

### Identification of mammalian GPCR homologs in fungal species

Functional Pfams from each mammalian GPCR family include 7tm 1 (Rhodopsin), 7tm 3 (Gluta-mate), Frizzled/Fz (Frizzled), TAS2R (Taste2), and GAIN/GPS/hEGF/EGF CA/Calx-beta/Cadher (Adhesion). These were identified in 1,357 fungal species, and hits with an p-value <1e-05 containing these Pfam domains were verified using TMHMM [49], retaining only sequences with 7 TMHs.

### Comparative phylogenetic analysis of Pth11-like GPCRs

Phylogenetic trees have been constructed for a specific clade of species within Pezizomycotina (Supplementary file 1 Table S6) that are closely related and exhibit substantial variations in the abundance of Class 14 GPCRs. Initially, we conducted a search for species of interest within the Pezizomycotina species tree available on the JGI genome portal [48]. Ultimately, we identified 12 species (Table 2) that share close evolutionary relationships. These species also had more than 10 times the abundance of Pth11-like GPCRs compared to other species. The species tree for the 12 genomes and the gene tree for Pth11-like GPCRs were built using Orthofinder version 2.5.4 [53]. Both trees were visualized and compared using iTOL version 6.9.1 [54].

### Comparative Gene families enrichment analysis

OrthoFinder was employed to identify orthogroups in species exhibiting substantial differences in Pth11-like GPCR abundance. Notung version 2.9.1 [34] was used to infer duplication and loss events for Pth11-like GPCRs. To detect expanded and contracted gene families, we applied CAFE5 [35], filtering significantly expanded or contracted families using a threshold of *p* < 0.05. Gene Ontology (GO) enrichment analysis was executed using TBtools-II [55] and Pfam domain enrichment analysis were performed on these gene families, and statistical significance was assessed using a two-sided test with p-values adjusted via the Benjamini-Hochberg method [56].

### Potential novel GPCR prediction

#### 1D-CCN architecture

We implemented a 1D Convolutional Neural Network (1D-CNN) to classify protein sequences as either GPCR or non-GPCR. Positive data consists of all 17,996 sequences from all 14 classes of GPCRs that we identified. For the negative dataset, we collected 18,221 sequences containing 7 TMHs, with Pfam domains associated with protein functions unrelated to GPCR activity (Table 1). The sequences were one-hot encoded, converting each amino acid into a binary vector, and padding was applied to manage variable sequence lengths. Then the data was split into 80% for training and 20% for testing. The model architecture consists of a 1D convolutional layer with 32 filters, a kernel size of 3, and ReLU activation. This is followed by a max-pooling layer with a pool size of 2, which reduces the dimensionality of the feature maps. The flattened output is passed through a fully connected layer with 64 neurons, also with ReLU activation [57], and a sigmoid-activated output layer for binary classification. The model was compiled using the Adam optimizer [58] and binary cross-entropy loss, and its performance was evaluated based on accuracy. After training for 10 epochs with a batch size of 32, the model achieved a test accuracy of 0.97.

#### Preparation of query sequences

First, we searched all 1,357 fungal genomes in the database for sequences with 7 TMHs using TMHMM [49]. 31,493 newly identified GPCRs were filtered out. Next, MCL clustering [59] was performed. For the largest cluster, multiple sequence alignment was conducted using MAFFT [50] to assess sequence similarity. Sequences from the largest subcluster containing the DUF6534 domain were analyzed to determine whether they represent a novel GPCR.

#### GPCR-G-alpha protein interaction prediction

Well-known GPCR and G-alpha pairs, along with non-interacting pairs, were modeled using Alphafold Multimer version 2.2 [33]. Potential novel GPCRs from 155 species were then paired with G-alpha proteins from the same species and processed through AlphaFold Multimer. The pairs with the highest confidence scores in each species were selected, and the corresponding.pdb and.pkl files were collected for pDockQ2 score calculation [43].

AlphaFold Multimer has been utilized to validate protein complexes, whereas methods such as pDockQ2 [43] evaluate interface quality through predicted aligned error (PAE) and predicted Local Distance Difference Test (pLDDT) [42] metrics in the absence of native structures.

## Acknowledgements

The work conducted by the U.S. The Department of Energy Joint Genome Institute (https://ror.org/04xm1d337), a DOE Office of Science User Facility, is supported by the Office of Science of the U.S. Department of Energy operated under Contract No. DE-AC02-05CH11231.

## References

[1] Daniel M Rosenbaum, Søren GF Rasmussen, and Brian K Kobilka. The structure and function of G-protein-coupled receptors. Nature, 459(7245):356–363, 2009.

[2] Raise Ahmad, Stefanie Wojciech, and Ralf Jockers. Hunting for the function of orphan GPCRs–beyond the search for the endogenous ligand. British Journal of Pharmacology, 172(13):3212–3228, 2015.

[3] Sabine Gruber, Markus Omann, and Susanne Zeilinger. Comparative analysis of the repertoire of G protein-coupled receptors of three species of the fungal genus Trichoderma. BMC Microbiology, 13:1–14, 2013.

[4] Fred Naider and Jeffrey M Becker. The *α*-factor mating pheromone of Saccharomyces cerevisiae: a model for studying the interaction of peptide hormones and G protein-coupled receptors. Peptides, 25(9):1441–1463, 2004.

[5] Yonglin Wang, Aining Li, Xiaoli Wang, Xin Zhang, Wei Zhao, Daolong Dou, Xiaobo Zheng, and Yuanchao Wang. GPR11, a putative seven-transmembrane G protein-coupled receptor, controls zoospore development and virulence of Phytophthora sojae. Eukaryotic Cell, 9(2):242–250, 2010.

[6] C Braunsdorf, D Mailänder-Sánchez, and M Schaller. Fungal sensing of host environment. Cellular Microbiology, 18(9):1188–1200, 2016.

[7] Margarita Juárez-Montiel, Daniel Clark-Flores, Pedro Tesillo-Moreno, Esaú de la Vega-Camarillo, Dulce Andrade-Pavón, Juan Alfredo Hernández-García, César Hernández-Rodríguez, and Lourdes Villa-Tanaca. Vacuolar proteases and autophagy in phytopathogenic fungi: A review. Frontiers in Fungal Biology, 3:948477, 2022.

[8] Yong-Sun Bahn, Chaoyang Xue, Alexander Idnurm, Julian C Rutherford, Joseph Heitman, and Maria E Cardenas. Sensing the environment: lessons from fungi. Nature Reviews Microbiology, 5(1):57–69, 2007.

[9] JF Martín, MA Van Den Berg, E Ver Loren Van Themaat, and P Liras. Sensing and transduction of nutritional and chemical signals in filamentous fungi: Impact on cell development and secondary metabolites biosynthesis. Biotechnology Advances, 37(6):107392, 2019.

[10] Jeong-Ah Seo, Kap-Hoon Han, and Jae-Hyuk Yu. The gprA and gprB genes encode putative G protein-coupled receptors required for self-fertilization in Aspergillus nidulans. Molecular Microbiology, 53(6):1611–1623, 2004.

[11] Yong Xue, Montserrat Batlle, and Jeanne P Hirsch. GPR1 encodes a putative G protein-coupled receptor that associates with the Gpa2p G*α* subunit and functions in a Ras-independent pathway. The EMBO Journal, 1998.

[12] Leon Kraakman, Katleen Lemaire, Pingsheng Ma, Aloys WRH Teunissen, Monica CV Donaton, Patrick Van Dijck, Joris Winderickx, Johannes H De Winde, and Johan M Thevelein. A Saccharomyces cerevisiae G-protein coupled receptor, Gpr1, is specifically required for glucose activation of the cAMP pathway during the transition to growth on glucose. Molecular Microbiology, 32(5):1002–1012, 1999.

[13] Kyung-Sook Chung, Misun Won, Sang-Bong Lee, Young-Joo Jang, Kwang-Lae Hoe, Dong-Uk Kim, Ji-Won Lee, Kyu-Won Kim, and Hyang-Sook Yoo. Isolation of a Novel Gene fromSchizosaccharomyces pombe: stm1+ Encoding a Seven-transmembrane Loop Protein That May Couple with the Heterotrimeric G*α*2 Protein, Gpa2. Journal of Biological Chemistry, 276(43):40190–40201, 2001.

[14] James E Galagan, Sarah E Calvo, Katherine A Borkovich, Eric U Selker, Nick D Read, David Jaffe, William FitzHugh, Li-Jun Ma, Serge Smirnov, Seth Purcell, et al. The genome sequence of the filamentous fungus Neurospora crassa. Nature, 422(6934):859–868, 2003.

[15] Mohamed MH El-Defrawy and Abd El-Latif Hesham. G-protein-coupled receptors in fungi. Fungal Biotechnology and Bioengineering, pages 37–126, 2020.

[16] Katherine A Borkovich, Lisa A Alex, Oded Yarden, Michael Freitag, Gloria E Turner, Nick D Read, Stephan Seiler, Deborah Bell-Pedersen, John Paietta, Nora Plesofsky, et al. Lessons from the genome sequence of Neurospora crassa: tracing the path from genomic blueprint to multicellular organism. Microbiology and Molecular Biology Reviews, 68(1):1–108, 2004.

[17] Anne Lafon, Kap-Hoon Han, Jeong-Ah Seo, Jae-Hyuk Yu, and Christophe d’Enfert. G-protein and cAMP-mediated signaling in aspergilli: a genomic perspective. Fungal Genetics and Biology, 43(7):490–502, 2006.

[18] Resham D Kulkarni, Michael R Thon, Huaqin Pan, and Ralph A Dean. Novel G-protein-coupled receptor-like proteins in the plant pathogenic fungus Magnaporthe grisea. Genome Biology, 6:1–14, 2005.

[19] Thomas J Lyons, Nancy Y Villa, Lisa M Regalla, Brian R Kupchak, Anna Vagstad, and David J Eide. Metalloregulation of yeast membrane steroid receptor homologs. Proceedings of the National Academy of Sciences U. S. A, 101(15):5506–5511, 2004.

[20] Manolis Kellis, Nick Patterson, Matthew Endrizzi, Bruce Birren, and Eric S Lander. Sequencing and comparison of yeast species to identify genes and regulatory elements. Nature, 423(6937):241–254, 2003.

[21] Christiam Camacho, George Coulouris, Vahram Avagyan, Ning Ma, Jason Papadopoulos, Kevin Bealer, and Thomas L Madden. BLAST+: architecture and applications. BMC Bioinformatics, 10:1–9, 2009.

[22] Hongxia Zheng, Lei Zhou, Tonghai Dou, Xiaotian Han, Yanyan Cai, Xiaoying Zhan, Cheng Tang, Jing Huang, and Qihan Wu. Genome-wide prediction of G protein-coupled receptors in Verticillium spp. Fungal Biology, 114(4):359–368, 2010.

[23] Ilva E Cabrera, Itallia V Pacentine, Andrew Lim, Nayeli Guerrero, Svetlana Krystofova, Liande Li, Alexander V Michkov, Jacqueline A Servin, Steven R Ahrendt, Alexander J Carrillo, et al. Global analysis of predicted G protein-coupled receptor genes in the filamentous fungus, Neurospora crassa. G3: Genes, Genomes, Genetics, 5(12):2729–2743, 2015.

[24] Helgi B Schiöth and Robert Fredriksson. The GRAFS classification system of G-protein coupled receptors in comparative perspective. General and Comparative Endocrinology, 142(1-2):94–101, 2005.

[25] Ravikrishna Ramanujam, Meredith E Calvert, Poonguzhali Selvaraj, and Naweed I Naqvi. The late endosomal HOPS complex anchors active G-protein signaling essential for pathogenesis in Magnaporthe oryzae. PLoS Pathogens, 9(8):e1003527, 2013.

[26] Liande Li, Sara J Wright, Svetlana Krystofova, Gyungsoon Park, and Katherine A Borkovich. Heterotrimeric G protein signaling in filamentous fungi. Annu. Rev. Microbiol., 61(1):423–452, 2007.

[27] Xihui Xu, Guopeng Li, Lu Li, Zhenzhu Su, and Chen Chen. Genome-wide comparative analysis of putative Pth11-related G protein-coupled receptors in fungi belonging to Pezizomycotina. BMC Microbiology, 17:1–11, 2017.

[28] Rosalin Bonetta and Gianluca Valentino. Machine learning techniques for protein function prediction. Proteins: Structure, Function, and Bioinformatics, 88(3):397–413, 2020.

[29] Divyanshu Aggarwal and Yasha Hasija. A review of deep learning techniques for protein function prediction. arXiv preprint arXiv:2211.09705, 2022.

[30] Theo Sanderson, Maxwell L Bileschi, David Belanger, and Lucy J Colwell. ProteInfer, deep neural networks for protein functional inference. Elife, 12:e80942, 2023.

[31] Vladimir Gligorijević, P Douglas Renfrew, Tomasz Kosciolek, Julia Koehler Leman, Daniel Berenberg, Tommi Vatanen, Chris Chandler, Bryn C Taylor, Ian M Fisk, Hera Vlamakis, et al. Structure-based protein function prediction using graph convolutional networks. Nature Communications, 12(1):3168, 2021.

[32] Samuel Sledzieski, Rohit Singh, Lenore Cowen, and Bonnie Berger. D-SCRIPT translates genome to phenome with sequence-based, structure-aware, genome-scale predictions of protein-protein interactions. Cell Systems, 12(10):969–982, 2021.

[33] Richard Evans, Michael O’Neill, Alexander Pritzel, Natasha Antropova, Andrew Senior, Tim Green, Augustin Žídek, Russ Bates, Sam Blackwell, Jason Yim, et al. Protein complex prediction with AlphaFold-Multimer. biorxiv, pages 2021–10, 2021.

[34] Maureen Stolzer, Katherine Siewert, Han Lai, Minli Xu, and Dannie Durand. Event inference in multidomain families with phylogenetic reconciliation. BMC Bioinformatics, 16:1–20, 2015.

[35] Tijl De Bie, Nello Cristianini, Jeffery P Demuth, and Matthew W Hahn. CAFE: a computational tool for the study of gene family evolution. Bioinformatics, 22(10):1269–1271, 2006.

[36] Igor R Pozdnyakov, Evgeniy V Potapenko, Elena S Nassonova, Vladislav V Babenko, Daria I Boldyreva, Victoria S Tcvetkova, and Sergey A Karpov. To the Origin of Fungi: Analysis of MFS Transporters of First Assembled Aphelidium Genome Highlights Dissimilarity of Osmotrophic Abilities between Aphelida and Fungi. Journal of Fungi, 9(10):1021, 2023.

[37] Ján Víglaš and Petra Olejníková. An update on ABC transporters of filamentous fungi–from physiological substrates to xenobiotics. Microbiological Research, 246:126684, 2021.

[38] Nils Thieme, Vincent W Wu, Axel Dietschmann, Asaf A Salamov, Mei Wang, Jenifer Johnson, Vasanth R Singan, Igor V Grigoriev, N Louise Glass, Chris R Somerville, et al. The transcription factor PDR-1 is a multi-functional regulator and key component of pectin deconstruction and catabolism in Neurospora crassa. Biotechnology for Biofuels, 10:1–21, 2017.

[39] Julia Koehler Leman, Pawel Szczerbiak, P Douglas Renfrew, Vladimir Gligorijevic, Daniel Berenberg, Tommi Vatanen, Bryn C Taylor, Chris Chandler, Stefan Janssen, Andras Pataki, et al. Sequence-structure-function relationships in the microbial protein universe. Nature Communications, 14(1):2351, 2023.

[40] Henrik G Dohlman, Jianping Song, Doreen Ma, William E Courchesne, and Jeremy Thorner. Sst2, a negative regulator of pheromone signaling in the yeast Saccharomyces cerevisiae: expression, localization, and genetic interaction and physical association with Gpa1 (the G-protein *α* subunit). Molecular and Cellular Biology, 16(9):5194–5209, 1996.

[41] Kendall J Blumer and Jeremy Thorner. Beta and gamma subunits of a yeast guanine nucleotide-binding protein are not essential for membrane association of the alpha subunit but are required for receptor coupling. Proceedings of the National Academy of Sciences U. S. A, 87(11):4363–4367, 1990.

[42] Valerio Mariani, Marco Biasini, Alessandro Barbato, and Torsten Schwede. lDDT: a local superposition-free score for comparing protein structures and models using distance difference tests. Bioinformatics, 29(21):2722–2728, 2013.

[43] Wensi Zhu, Aditi Shenoy, Petras Kundrotas, and Arne Elofsson. Evaluation of AlphaFold-Multimer prediction on multi-chain protein complexes. Bioinformatics, 39(7):btad424, 2023.

[44] Chengshu Wang and Sibao Wang. Insect pathogenic fungi: genomics, molecular interactions, and genetic improvements. Annual Review of Entomology, 62(1):73–90, 2017.

[45] Rony Chamoun, Konstantinos A Aliferis, and Suha Jabaji. Identification of signatory secondary metabolites during mycoparasitism of Rhizoctonia solani by Stachybotrys elegans. Frontiers in Microbiology, 6:353, 2015.

[46] Tom JB de Man, Jason E Stajich, Christian P Kubicek, Clotilde Teiling, Komal Chenthamara, Lea Atanasova, Irina S Druzhinina, Natasha Levenkova, Stephanie SL Birnbaum, Seth M Barribeau, et al. Small genome of the fungus Escovopsis weberi, a specialized disease agent of ant agriculture. Proceedings of the National Academy of Sciences U. S. A., 113(13):3567– 3572, 2016.

[47] Lotus A Lofgren, Nhu H Nguyen, Rytas Vilgalys, Joske Ruytinx, Hui-Ling Liao, Sara Branco, Alan Kuo, Kurt LaButti, Anna Lipzen, William Andreopoulos, et al. Comparative genomics reveals dynamic genome evolution in host specialist ectomycorrhizal fungi. New Phytologist, 230(2):774–792, 2021.

[48] Igor V Grigoriev, Roman Nikitin, Sajeet Haridas, Alan Kuo, Robin Ohm, Robert Otillar, Robert Riley, Asaf Salamov, Xueling Zhao, Frank Korzeniewski, et al. MycoCosm portal: gearing up for 1000 fungal genomes. Nucleic Acids Research, 42(D1):D699–D704, 2014.

[49] Anders Krogh, Björn Larsson, Gunnar Von Heijne, and Erik LL Sonnhammer. Predicting transmembrane protein topology with a hidden Markov model: application to complete genomes. Journal of Molecular Biology, 305(3):567–580, 2001.

[50] Kazutaka Katoh, Kazuharu Misawa, Kei-ichi Kuma, and Takashi Miyata. MAFFT: a novel method for rapid multiple sequence alignment based on fast Fourier transform. Nucleic Acids Research, 30(14):3059–3066, 2002.

[51] Salvador Capella-Gutiérrez, José M Silla-Martínez, and Toni Gabaldón. trimAl: a tool for automated alignment trimming in large-scale phylogenetic analyses. Bioinformatics, 25(15):1972–1973, 2009.

[52] Sean R Eddy. Accelerated profile HMM searches. PLoS Computational Biology, 7(10):e1002195, 2011.

[53] David M Emms and Steven Kelly. OrthoFinder: phylogenetic orthology inference for comparative genomics. Genome Biology, 20:1–14, 2019.

[54] Ivica Letunic and Peer Bork. Interactive Tree of Life (iTOL) v6: recent updates to the phylogenetic tree display and annotation tool. Nucleic Acids Research, page gkae268, 2024.

[55] Chengjie Chen, Ya Wu, Jiawei Li, Xiao Wang, Zaohai Zeng, Jing Xu, Yuanlong Liu, Junting Feng, Hao Chen, Yehua He, et al. TBtools-II: A “one for all, all for one” bioinformatics platform for biological big-data mining. Molecular Plant, 16(11):1733–1742, 2023.

[56] Yoav Benjamini and Yosef Hochberg. Controlling the false discovery rate: a practical and powerful approach to multiple testing. Journal of the Royal Statistical Society: Series B (Methodological), 57(1):289–300, 1995.

[57] AF Agarap. Deep learning using rectified linear units (relu). arXiv preprint arXiv:1803.08375, 2018.

[58] Diederik P Kingma. Adam: A method for stochastic optimization. arXiv preprint arXiv:1412.6980, 2014.

[59] Stijn Van Dongen. Graph clustering via a discrete uncoupling process. SIAM Journal on Matrix Analysis and Applications, 30(1):121–141, 2008.

